# A heuristic feature cluster search algorithm for precise functional brain mapping

**DOI:** 10.1101/518480

**Authors:** Nima Asadi, Yin Wang, Ingrid Olson, Zoran Obradovic

## Abstract

Detecting the most relevant brain regions for explaining the distinction between cognitive conditions is one of the most sought after objectives in neuroimaging research. A popular approach for achieving this goal is the multivariate pattern analysis (MVPA) which is commonly conducted through the searchlight procedure as well as a number of other approaches. This is due to advantages of such methods which include being intuitive and flexible with regards to size of the search space. However, these approaches suffer from a number of limitations that lead to misidentification of truly informative voxels or clusters of voxels which in turn results in imprecise information maps. The limitations of such procedures mainly stem from several factors such as the fact that the information value of the search spheres are assigned to the voxel at the center of them (in case of searchlight), the requirement for manual tuning of parameters such as searchlight radius and shape and other optimization parameters, overlooking the structure and interactions within the regions, and the drawbacks of using regularization methods in analysis of datasets with characteristics of common fMRI data. In this paper, we propose a fully data-driven maximum relevance minimum redundancy search algorithm for detecting precise information value of voxel-level clusters within brain regions while alleviating the above mentioned limitations. In order to make the algorithm efficient, we propose an implementation based on principles of dynamic programming. We evaluate and compare the proposed algorithm with the searchlight procedure using both real and synthetic datasets.

## Introduction

In the common form of functional magnetic resonance imaging (fMRI), the blood-oxygen level dependent (BOLD) contrast is extracted as the response signal in order to measure neural activity in the brain^1^. Measurement of this response signal over time forms a time course corresponding to each voxel whose dimensions depend on the spatial resolution of the imaging device. Analysis and comparison of these time courses can reveal valuable knowledge regarding different neurological conditions among populations. Popular approaches for analyzing fMRI data can be broken down into two main categories: voxel-wise univariate analysis, and multi-voxel pattern analysis, also known as MVPA^2, 3^. The univariate analysis searches for correlations between psychological or physical status and the activation of single voxels while MVPA aims to detect patterns among conditions observed among combinations of multiple voxels ^4^. Unlike univariate analyses, MVPA approaches are designed to allow researchers to test how dispersed patterns of BOLD activation across multiple voxels relate to experimental conditions^4, 5^. One approach in multi-voxel scheme is to compare and analyze spatially averaged (smoothed) BOLD activations across the entire regions of interest. Advantages of this approach include an increase in the signal to noise ratio as well as the consistency of the analysis among subjects can be noted ^6^. However, spatial smoothing leads to significant loss of information about the patterns of activation within the regions of interest. This information includes the activities and dynamics within subregions which can provide valuable insight into their relation with different mental states^7, 8^. This issue becomes more complex when dealing with larger regions of interest. Therefore, in order to capture such information, it is necessary to consider the BOLD activity in smaller spherical subsets^9^.

The question of identifying relevant regions with regards to specific conditions has prompted numerous studies during the recent decades. One of the most commonly employed approaches for this application is the searchlight method proposed by Kriegeskorte et al., which given the dimensions of a sphere window, performs a search across a brain region to detect the information of sets of neighboring voxels^10, 11^. In this multivariate approach, spatial patterns of activity within the search window are compared between two groups using statistical discriminant analysis or supervised machine learning approaches ^12, 13^. The search sphere (“searchlight”) is centered on every voxel, i.e. the derived separability value for each voxel is derived from the discrimination score of its surrounding searchlight, not the voxel individually. Advantages of searchlight analysis include its data-driven nature, its ability in performing whole-brain search without the need to specify brain regions, and its high interpretability.

However, the searchlight procedure suffers from critical drawbacks which can lead to erroneous detection of informative voxels/regions. Etzel et al. discussed several issues with the searchlight method in detail which we briefly point out here^9^. One limitation of the searchlight procedure is that it can declare a subregion with a few highly-informative voxels as informative, making detection of informative voxel clusters ambiguous. This issue becomes more prevalent with selection of larger search radii^9^. Moreover, choosing an appropriate search radius is essential, which depends on the shape and size of the region being searched. However, finding the discriminative subregion by a search over several possible search radius values is difficult specially when being applied to whole-brain analysis. Aside from this issue, the shape of the searchlight can limit the detection of the subregions with the highest discrimination power. This is due to the fact that the searchlight is commonly in the shape of a sphere or a cube, which forces subregions with irregular shapes to fall between multiple searchlight positions. This issue can partially be relieved through assigning the searchlight sphere as small as possible at the expense of overfitting^9^. Another shortcoming with this method is the fact that assignment of a single searchlight radius might provide optimal results for one subregion, but does not guarantee similar results for many other regions. Consequently, finding the optimal searchlight radius for a large search space comprised of subspaces with varying anatomical characteristics is a challenging task. Tackling some of the mentioned issues requires further analysis while some issues are inherently irresolvable through the scope of the searchlight procedure.

Several other approaches have been proposed based on machine learning techniques to create models for automated decoding of cognitive states during recent years. A number of these techniques proposed using different variations of the least absolute shrinkage and selection operator(Lasso) family to develop a continuous feature evaluation criteria^14–17^. However, the use of lasso regularization in fMRI studies introduces several limitations. One of such constraints is the fact that in case of number of features *p* being larger than the number of examples *m*, lasso selects *m* features at maximum^18^. This is a critical drawback due to the fact that in fMRI studies, specially on voxel-level analysis, it is very common that the number of subjects is far smaller than the number of features (voxels, or even regions of interest). Another drawback of Lasso is the fact that since it forces less important coefficients to be zero, it does not provide the information value of the features that have not been selected. Consequently, instead of creating an information spectrum, it points to a small subset of features that it finds to be more informative, which makes it less useful for researchers in fMRI studies due to loss of knowledge regarding majority of the brain areas. Moreover, using a more recent variation of Lasso which considers group structure named group Lasso requires disjoint subsets of the voxels to be pre-determined. This limitation creates an issue similar to the searchlight analysis radius selection since the choice of size and structure of the groups of voxels changes the results of the feature space shrinkage. Also, the interpretability of performing a regularization-based approach on the entire feature space is low. Another method for detecting biomarkers is the manifold learning suggested by ^19^. Despite its power in nonlinear classification of MR images and the consideration of spectral theory in dimensionality reduction, several parameters need to be fine tuned for it to achieve preferable results. These parameters include the optimal neighborhood size, the number of dimensions learned by the manifold, and the heat kernel parameter which the Laplacian eigenmap feature selection is sensitive to. Also, time complexity of the spectral embedding phase of manifold learning grows substantially with the size of neighborhood, making it less efficient for full-brain analysis^20^.

Therefore, development of new analytical models is essential for the critical task of automatically discovering the information of different regions regarding certain neurological conditions to tackle the above-mentioned issues. In this study, we propose a new algorithm for discovering the discriminant power of different brain regions for explaining the disparities in neural activities among populations. The goal of this work is to provide a mapping of information clusters where the spatial proximity, structural characteristics of the voxels, and the gradient neural activation patterns are taken into account without the requirement of parameter tuning. Through a completely data-driven search, the proposed approach achieves this goal while increasing the precision of information cluster discovery and classification accuracy at the same time. Through empirical results on a real fMRI dataset as well as synthetic data, we compare the performance of the proposed algorithm with the searchlight methods. We explain the experimental results as well as the suggested methodology in more detail in the next sections.

The objective of the proposed methodology is to create the information map of the brain (or regions of interest) with regards to a certain neurological status, e.g. a cognitive disorder, age, activation differences between tasks, etc. In other words, given two (or more) populations, the goal is to discover the level at which the BOLD activation of brain regions differ between groups, which in this study we defined as the discriminability score of the brain region (Note that we use the terms discriminability score and information interchangeably in order to preserve consistency with the related literature). In other words, the discriminability score is the relevance of a feature (the neural activity or corresponding BOLD value of a voxel) or a set of features (neural activity of a cluster of voxels) in separating two or more classes of subjects. Note that clusters of voxels are groups of neighboring voxels whose size can span from one voxel to the entire region of interest. We also define a search space as the region of brain that we intend to explore (search) and investigate in order to detect informative subregions. Generally, the input to the proposed approach is a set of matrices, each belonging to a subject, where each element in the matrices is the activation or BOLD value of a voxel in the search space averaged over time. The output of the proposed algorithm is an information map of the search space where each voxel/cluster is assigned a discriminability score. In order to simplify reference to the proposed algorithm, we use the term MNS, which is the acronyms for maximum relevance minimum redundancy (MRMR) neighborhood search.

### The Proposed Algorithm

To extract the informative clusters within a region of interest we propose an algorithm which traverses the search space based on a information-based heuristic, and outputs the discovered information clusters and their measured discriminant scores after termination. This criteria is similar to greedy search algorithms which pick the “best” neighbor according to a heuristic, which in this case is the discriminability score of the cluster of voxels^21, 22^. However, unlike common greedy algorithm procedures, the proposed algorithm reviews the searched clusters and prunes the redundant voxels after each expansion step. A pseudo code of the proposed algorithm is provided in Algorithm 1 in the methodology section along with implementation details. The MNS feature selection algorithm is loosely inspired by path greedy finding algorithms such as A* search, and variable neighborhood search strategies^23–27^. However, in order to formulate our problem into a spatial search problem, we propose several modifications and constraints: 1-We use the spectral feature quality as our heuristic,2- instead of setting goal states as a node that the algorithm needs to reach, we set it as maximum discriminant score possible in the accessible neighborhood, 3- the search at each step is performed on the neighbors of the combination of chosen voxels (information cluster) rather than on individual voxels, 4-and unlike the mentioned search algorithms, the proposed algorithm does not necessarily meet every voxel in the search space, but instead, it terminates when there is no more admissible voxels in the clusters neighborhood. In general, MNS performs a search starting from a voxel *V* to detect its immediate neighbors whose pairing with *V* enhances its power in distinguishing between the classes of data examples. This criteria is similar to the maximum relevance criteria in feature selection. Then, a redundancy detection is performed on the newly created cluster to remove the redundant features and further optimize the selected voxels (features). After this redundancy procedure, similar relevance criteria is performed on the remaining cluster, meaning that the neighboring voxels of the entire cluster are examined to find the useful voxels to add to the cluster and expand it. This criteria is performed until there are no neighboring voxels left whose addition to the detected cluster is helpful. In other words, the proposed search procedure relaxes the requirement of performing multiple searches starting from a certain voxel while increasing the spatial precision of search by investigating voxel-level resolutions. Searching the neighborhood of the entire clusters is advantageous for considering new formations and structures between groups of voxel in order to increase the generalizability of the results. Moreover, it alleviates the issue of local optima in greedy algorithms ^28, 28, 29^. The proposed approach and its analytical and computational details will be discussed in the next sections. In general, the steps of the proposed algorithm go as followed (details are provided in algorithm 1):

**Step 1** Start from voxel *v_s_* and measure the relevance score of its conjugation with each of its immediate neighbors one by one, and select the neighbors whose addition to *v_s_* increases its information. Then combine *v_s_* and its set of useful neighbors *V_u_* to create the information cluster *C_inf_*.
**Step 2** Search each voxel adjacent to *C_inf_*, and admit the neighboring voxels whose admission to *C_inf_* enhances its relevance.
**Step 3** Perform the information redundancy analysis on *C_inf_* and remove its redundant voxels with a bias against the margins of *C_inf_* cluster. Then move to the next neighborhood layer of the pruned *C_inf_* cluster.
**Step 4** Repeat Steps 2 and 3 and expand the cluster until there is no new neighbor whose addition to *C_inf_* increases its score. Save *C_inf_* and its information in the output variable.
**Step 5** Start from the voxel next to *v_s_* and follow steps 1 to 5.
**Step 6** When steps 1 to 4 are performed for every voxel as the starting voxel in the search space, terminate the algorithm and output the set of discovered information clusters and their information value.

As mentioned previously, the cluster originating from *V* is expanded through this criteria until no neighbors are found whose addition to the cluster enhances its discriminability score. In that case, the algorithm saves the detected cluster as well is its calculated score (information) as part of the output, and starts the same criteria starting from the voxel next to *V*. The algorithm terminates when a cluster is detected starting from every voxel in the search space. Note that the size of information clusters can span from one voxel (meaning that none of its immediate neighbors increase its information) to the entire search space (meaning that the entire search space as one cluster contains relevant information). However, both of these extreme cases were rare according to our experiments. This procedure is discussed in more detail in the material and methods section. Also, note that during step 2, rather than selecting only one neighbor, which is the criteria in many greedy search methods, a group of candidate voxels of each neighborhood layer of cluster *C_inf_* are admitted.

**Figure 1.**
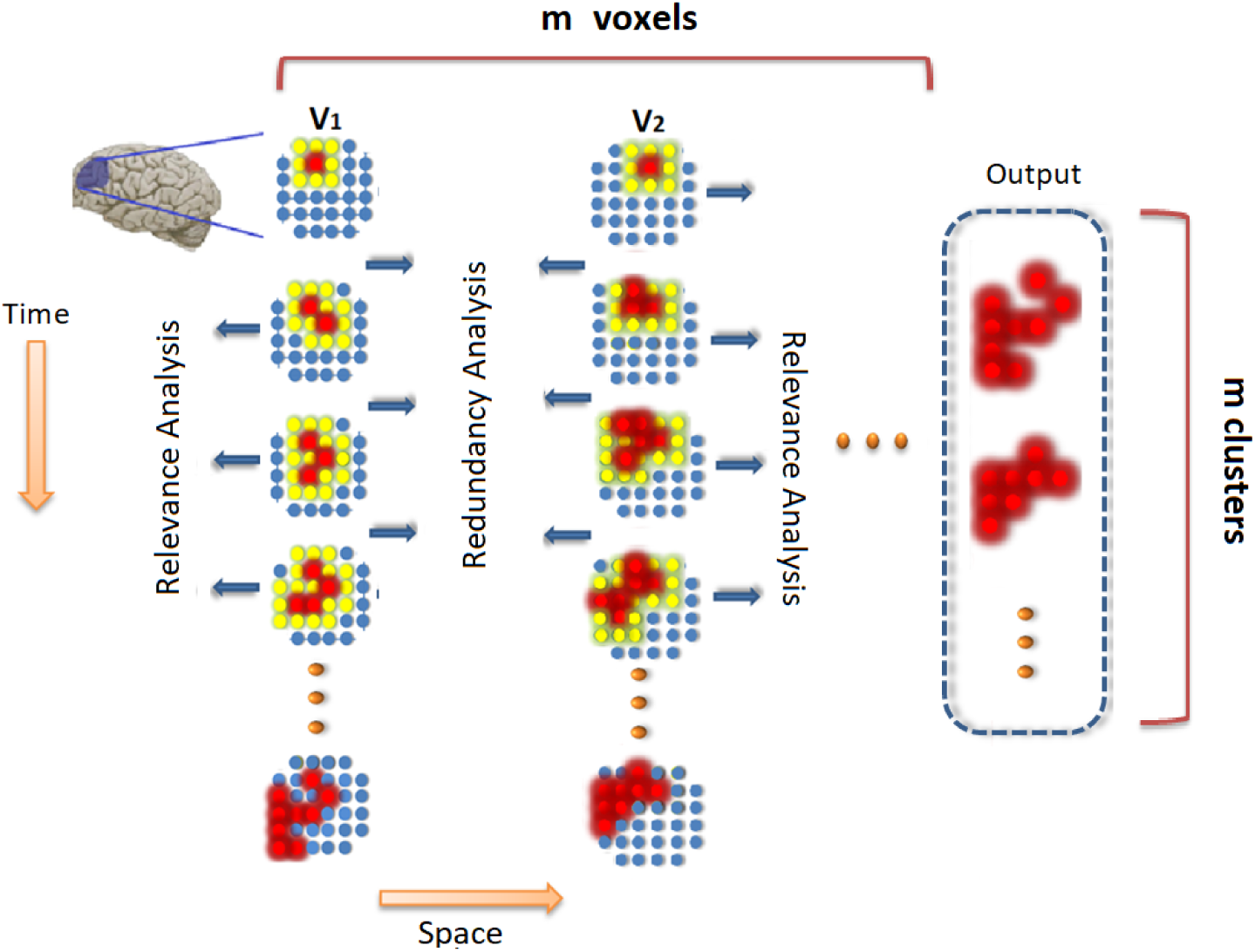
A schematic plot of of an example region traversed by the proposed algorithm The to detect information clusters originating from voxel *V*_1_ and *V*_2_. The algorithm first creates the information map for the cluster starting from *V*_1_ from top to bottom, and then moves to *V*_2_ and follows the same criteria. The red voxels are the admitted voxels and the yellow voxels represent the neighbors of the cluster at each step. The online spectral relevance analysis is performed during each search in the neighborhood layer, and the group biased mutual information redundancy anaylsis is performed before each next search in the neighborhood layer.

While the proposed algorithm provides a general information-based framework for information detection, the proper choice of heuristic function for analysis of relevance and redundancy needs several considerations: first, it is important to take into account the interaction and structure within groups of voxels rather than considering them as merely a number of of voxels in a group. Second, feasibility of the analysis should be considered as the number of voxels in the search space can be too large for many feature selection methods due to their time complexity. For the first point, we propose an online feature selection criteria which takes the interaction between features in to account. For the second issue, we propose an algorithmic technique for implementation of the proposed method known as dynamic programming, which makes the analysis time-efficient for experimentation on bigger search spaces such as whole brain analysis. In the next section, we describe these methodological techniques.

### Online Feature Selection as Heuristic Function

A basic approach for evaluating the discriminant power of a group of feature is the supervised feature subset selection by simply using the test accuracy of a trained machine learning algorithm. The major shortcomings of this approach include the requirement of retraining the model each time a new voxels information is being evaluated and its dependency on a specific model. Several other approaches have been proposed for feature set evaluation including statistical methods, linear discriminant analysis (LDA)^30, 31^ and spectral cluster analysis^32–34^. The purpose of all of these approaches is to provide a measure of how separable different groups of data are based on a feature group. These approaches are designed for offline feature selection where the entire feature group is known a priori. As explained previously, in the proposed MNS approach, the features flow into the model one at a time dynamically while the samples (subjects) are constant, and are added to the model if they are found to be beneficial to the information of the data, otherwise they are rejected. As a result, we can exploit this characteristic to formulate our search procedure as a criteria known as online (or streaming) feature selection, where evaluation of the features is performed by their arrival. Online feature selection is a relatively new topic where the number of observations is fixed but new features are added dynamically. During the recent years, a number of approaches have been proposed for online feature selection including the gradient descent based model named Grafting^35^, the likelihood ratio-based method named Alpha-investing^36^, and the feature redundancy and relevancy based method known as OSFS^37^. However, despite their advantages in feature selection, they do not capture the structure and correlation within the groups of voxels. Moreover,^38^ suggested an online group feature selection (OGFS), where the group structure of the features is considered in selecting the best subset. While this is a valuable quality, the use of Lasso in feature group analysis of their approach has limited capability for the domain of voxel level decoding of cognitive states due to previously mentioned issues. We incorporate an online feature selection method inspired by OSFS and OGFS as the heuristic function of our greedy algorithm. However, we use a different criteria for feature redundancy analysis while considering their interaction between the voxels and group structure as well as the gradient neural activation pattern within clusters. ere we explain the proposed heuristics for the suggested search approach.

**Figure 2.**
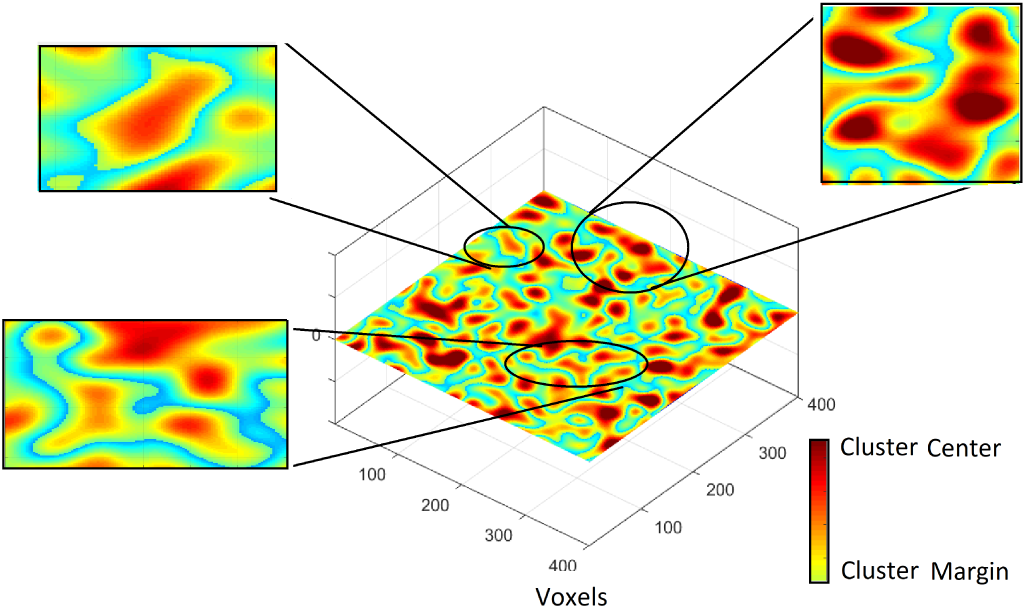
An example information map where the degree centrality of voxels within the cluster is shown via a heatmap. The voxels at the margins of the cluster have a lower degree centrality.

**Figure 3.**
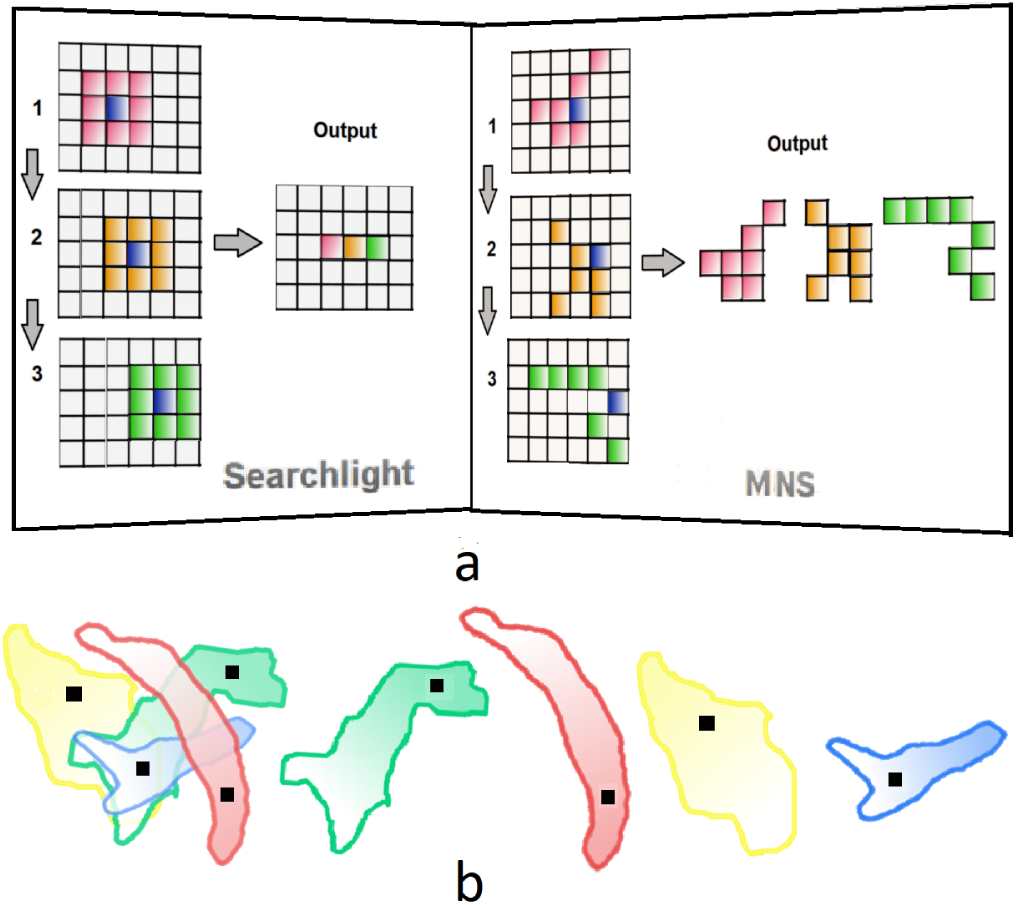
a.An schematic illustration of the steps and output of searchlight procedure compared with the MNS algorith. The blue voxel is the central voxel in the searchlight method, and the starting voxel in MNS algotihm. The radius for searchlight in this illustration is one voxel, and the information of the sphere, denoted by a specific color is assigned to the voxel at the center of the sphere, i.e. each voxel in the output map has the same color as its neighborhood. The output of the MNS method is a set of clusters which expanded from the starting voxel through a data-driven heuristic. The information of each clusters is demonstrated by a specific color. b.Left: An example illustration of overlapping clusters created by the proposed method. Right: the same clusters depicted individually. The voxel indicated by black dots are the starting voxel Vs which are expanded based on the discriminant analysis heuristic, resulting in a specific discriminant score for each cluster.

### Relevance Analysis

In order to form the information clusters, informative voxels are admitted based on relevance analysis criteria. This can be shown as the formula below:

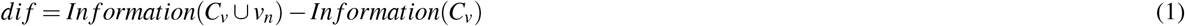

In other words, if the calculated discriminant power of the addition of the newly arrived voxel *v_n_* to cluster *C_v_*, is higher than the information of *C_v_* (*dif* > 0), the algorithm admits *v_n_* to the cluster *C_v_*. As the steps if the algorithm show, this relevance analysis is performed over all of the immediate neighbors of *C_v_*, and the relevant subset of the neighbors joins the cluster to expand it.

As mentioned before, several approaches are used for discriminant analysis of features. Note that many feature selection methods are designed for individual feature relevance analysis, however, in our application, the objective is to calculate the discriminant power of group of features (feature set selection). for this purpose, we used spectral feature analysis, particularly, group-level trace ratio of between-class (global) to within-class (local) affinity relationship in data^39^. This measure ensures that samples from the same class have a higher similarity compared with samples from different classes. Therefore, if addition of a new feature (dimension) increases this separability measure, it is considered as an admissible feature. A main reason for using this measure is its ability to consider global group information in admission of a new feature to the group. This measure is explained in more detail in the methodology section.

### Redundancy Analysis

Addition of new features to the group of features *C_v_* can introduce new redundancies among the existing features. In other words, the Markov blanket (MB) of the target variable defined as the optimal set of attributes to predict it can change due to the influence of the newly added feature^40, 41^. Therefore, by performing an inter-group feature subset shrinking before further expanding the cluster *C_v_* ∪ *v_n_*, we can find the optimal group of voxels *V_opt_* ∈ {*C_v_* ∪ *v_n_*}. This criteria further increases the precision of the information map by removing residual redundancies and increases the smoothness of the information clusters.

In order to perform this inter-group feature selection analysis, we not only consider the the mutual information between the set of features and the class variable but also the interaction between the features within the cluster. Through this approach, we were able capture more complex structures among groups of voxels rather than looking at their individual predictive power. In other words, the redundancy score of each feature is calculated as its own mutual information, subtract the mutual information between itself and the rest of the features in the group, plus the unconditional class-conditional correlations^42^. Therefore, there is an inverse relationship between the total value of this combination and the redundancy of the attribute. Moreover, in order to alleviate the noisy information existing in voxel-level resolution and consider the gradient proximal structure of neural activation, the voxels at the margins of the information clusters are penalized more compared to the voxel at the centers of the cluster. For this purpose, we used the neighborhood network within the clusters, and introduced a bias based on the degree centrality measure of the voxels within the cluster. Through this approach, a fast continuous redundancy analysis is created which is applied to the the voxels who have two characteristics: redundant to the prediction of the cognitive state, and their connection with the rest of the cluster is weak. This criteria helps us obtain smooth information clusters with minimal redundancy while taking to account the gradient pattern of activation in different brain regions. In figure 2 we can see example clusters created through the proposed algorithm in which the centrality areas within clusters are demonstrates by a heatmap. The details of both the relevance analysis and biased inter-group redundancy analysis is provided in the methodology section.

### Dynamic Programming Implementation

As shown in the previous section, the proposed algorithm performs a set of calculations in every step of the search to investigate every voxel at the vicinity of the detected clusters. Therefore, repetition of a large portion of the calculations occurs due to the search on overlapping regions. This significantly affects the analysis time and can make search over large regions infeasible. Dynamic Programming is a technique for making problems of recursive nature more efficient when the computations of the subproblems overlap^43, 44^. We use this approach to speed up both relevance and redundancy criteria by breaking down the calculations for every voxel and storing them at look up hash tables to be later used in the search process. At every step of the search, the algorithm first checks if the fragment of calculation encountered during the search already exists in the tables. If so, the algorithm directly uses the previously calculated values to save time, otherwise it performs the necessary calculation and stores it in the table for future use. The details of the dynamic programming implementation is provided in material and methods section. This approach significantly increases the search time, making precise information detection on voxel-level resolutions over large search spaces possible. Time complexity of MNS based on our experimentation setup is discussed in the results section.

### Interpretation of the Output

Due to data-driven expansion of the clusters by MNS criteria, emergence of overlapping clusters with different starting points is a possibility. Therefore, the output of the proposed method can be presented as a set of clusters with their measured discriminant scores instead of an information map. Figure 3 presents a comparison between the outputs of the searchlight procedure with the MNS algorithm after three steps of each method. As can be seen in that figure, the output of the searchlight algorithm is a map where the value of each voxel represents the information of the searchlight surrounding it while the output of MNS is a set of clusters originated from each voxel. In other words, the information of a cluster in MNS criteria is automatically assigned to the entire cluster. The same figure also shows how clusters originating from different voxels can overlap one another. The separated clusters on the right show a representation of the actual output where each cluster bears a certain information value. Therefore, each cluster is assigned an information score, and each voxel can be associated with multiple clusters. In machine learning point of view, each cluster of voxels can be considered a feature whose information value represents its classification quality. Therefore, a cluster with high information value is more suitable for prediction and diagnosis purposes.

### Experimental Setup and Data

The source code of the proposed method is available at https://github.com/ThisIsNima/GNS. All the experiments were performed on an Intel Core i7-3370 CPU, 3.40 GHz with 32 GB of RAM.

In order to evaluate the proposed approach, we first derived the information map of the fMRI datasets of two case studies based on the suggested algorithm as well as the searchlight procedure as the base line, and then selected the clusters which provide above chance (bigger than 50%) information as the classification features. Then, We compared the area under the curve (AUC) of a classifier which was trained on the two generated feature vectors to assess the quality of the generated features on the same dataset. For this purpose, we create two case studies using real fMRI data as well as a synthetic dataset.

The first case study includes a real fMRI dataset of 683 subjects from the publicly available Autism Brain Imaging Data Exchange (ABIDE) database^45^. This world-wide multi-site database includes resting state fMRI images of 370 healthy subjects, and 313 subjects diagnosed with Autism spectrum disorder (ASD). Also, despite the variances existing in this dataset due to diversity of data sources, we performed the analysis on the subjects as one group of data. Previously preprocessed rs-fMRI data was downloaded from the ABIDE database. This dataset was selected from the C-PAC preprocessing pipeline. The fMRI data was slice time corrected, motion corrected, and the voxel intensity was normalized using band-pass filtering and global signal regression. As mentioned in the methodology section, the input to the algorithm is the set of activation time courses of every voxel averaged over time, i.e. and *M* × *N* matrix where *M* is the number of subjects and *N* is the number of voxels in the search space.

To present the experimental results, we first compare the whole brain analysis performance, and for further investigation, we assign the search space to be regions of interest which are widely believed to play a crucial roles in ASD, namely the Hippocampus, Amygdalas, and Cerrebellums^46–49^. The Automated Anatomical Labeling (AAL) atlas was used to extract the regions of interest.

The second case study included simulated data where time courses of an average fMRI data with two conditions were generated for a population of 1000 subjects based on values extracted independently from a Gaussian distribution for four different sizes of search spaces of size 100 voxels, 500 voxels, 10,000 voxels, and 30,000 voxels. Also, noisy values were added to the signal with the constraints of realistic degree of correlation between adjacent voxels. A spatial pattern of response was then introduced to the two conditions which faded in and out according to the average temporal pattern of the cardiovascular response pattern among adults.

### Prediction results

The prediction results of the proposed approach is provided in Figures 6 and 7 where the area under the Receiver operating characteristic (ROC) curve is used as the evaluation measure.

After the discriminant scores of the clusters are revealed, the clusters with above chance information were used for classification. Note that based on both the MNS approach and the searchlight criteria a cluster was created for each voxel where in MNS, the cluster was expanded from each voxel *v_i_*, and for the searchlight procedure, it was the searchlight that encompassed each voxel *v_i_* by a certain radius. Similar spectral discriminant score of the relevance analysis in MNS was used as the analytical measure for the searchlight approach to make an appropriate comparison between the two approaches. In other words, the spectral discriminant score of each group of voxels surrounded by a search sphere was assigned to the voxel at the center of it. Based on both methods, each cluster contains an information value. The data was split in to train and test segments where 80% of the data was used for training, and 20% for testing. The information clusters were extracted from the training set. The clusters with over 50% discriminability power were then selected from the outputs of both methods, and two feature sets were created. Then, an SVM model was trained based on each of the feature sets, and was used to predict the labels of the test data. The reason for using a classification model (SVM) different than the information mapping models is to validate the generalizability of the analysis. par

#### Full brain analysis results

A visualization of the informative clusters (above 50% information) detected by MNS as well as the searchlight analysis is presented in Figure 4. As can be seen in that figure, the above chance clusters presented by the searchlight procedure show a lower discriminant score(maximum: 57.7%, average: 53.5%) compared to the MNS clusters (maximum: 75.1 %, average: 69.8 %). Also, while there are a number of regions where MNS and the searchlight detect similar informative areas, MNS detects an arrangement of voxels with a group structure which shows an increase in the information. These regions include the posterior cingulate cortex, Wernickarea, the amygdala, and left insula. Furthermore, MNS detected clusters with above chance information that the searchlight was not able to detect (their detected information were below 50%). These regions include portions of the cerebellum, and the anterior cingulate cortex.

Classification AUC of the two approaches using top features was also measured after an SVM model was trained based on both feature vectors separately. The search space for this analysis is the entire brain. For the searchlight method, radius values from 1 to 10 voxels were examined, and the highest performance, which belonged to the search sphere of 3 voxels, was selected. As can be seen in that plot, the classifier trained on the the MNS features on significantly outperforms the classifier trained on the searchlight features.

The precision of the two information decoding approaches on synthetic data is provided in Figure7 where the generated datasets contain 100, 500, 10,000, and 30,000 voxels. Similar to the real dataset, 5 fold train-test validation was also arranged for this set of experiments. Therefore, the data for 800 subjects were used for information mapping, and the remaining 200 subjects were used for classification test based on the top informative features according to the analysis on the training set. As can be seen in Figure7, the MNS method improves the classification accuracy over the searchlight procedure in all four setups.

#### Cluster level analysis results

The classification performance of the two approaches can also be assessed in smaller search spaces with more specific topological properties. This analysis is important due to the fact that assignment of a fixed searchlight radius on large search spaces might guarantee excellent performance in specific regions while underperform in other regions. In Figure 5, the classification accuracies based on every feature within the left crus II of the cerebellum (Region 93 per ATL) are compared between MNS and the searchlight criteria as an example. Analytical results of more regions are provided in the supplementary information. In that figure, the AUC value assigned to every voxel for the searchlight method is the calculated AUC that its search neighborhood with 3 voxel radius provides, i.e. the information of the individual clusters (searchlights). For MNS, this value for every voxel corresponds to the measured AUC for the cluster originated from it. Five runs of analysis based on both methods are presented in that figure, and due to the fluctuations in the AUC values as a result of random selection of the train-test samples, the average AUCs are indicated by the black line. As can be seen in that figure, the MNS method consistently demonstrates superiority in terms of classification accuracy compared to the searchlight approach. More comparisons of the two mapping methods on specific regions of interest are provide in Figure6, which further displays improvement of information map accuracy from the MNS method.

The experimental examinations of the proposed information mapping algorithm shows a significant enhancement over the searchlight procedure over both real and synthetic datasets. The higher classification test accuracy of MNS points to the data-driven advantage of identifying the informative combination of voxels that form various informative clusters. This is due to the fact that besides considering the combination of the voxels in their proximity, the formation structure of the groups of voxels as well as their interaction with one another also play important role in the information they provide. Moreover, redundant voxels are dynamically removed by MNS, which contributes to further enhance the precision of the discovery. These qualities are assessed through the online spectral feature evaluation and the spatially biased mutual information procedure. In other words, the MNS increases the information cluster dynamically as it searches for informative voxels to recruit for expanding the cluster while reassessing the Markov blanket in the existing feature set and removing the redundancies. While high precision is a crucial quality for information mapping of the brain, the proposed procedure benefits from other advantageous characteristics which are discussed in the next section.

**Figure 4.**
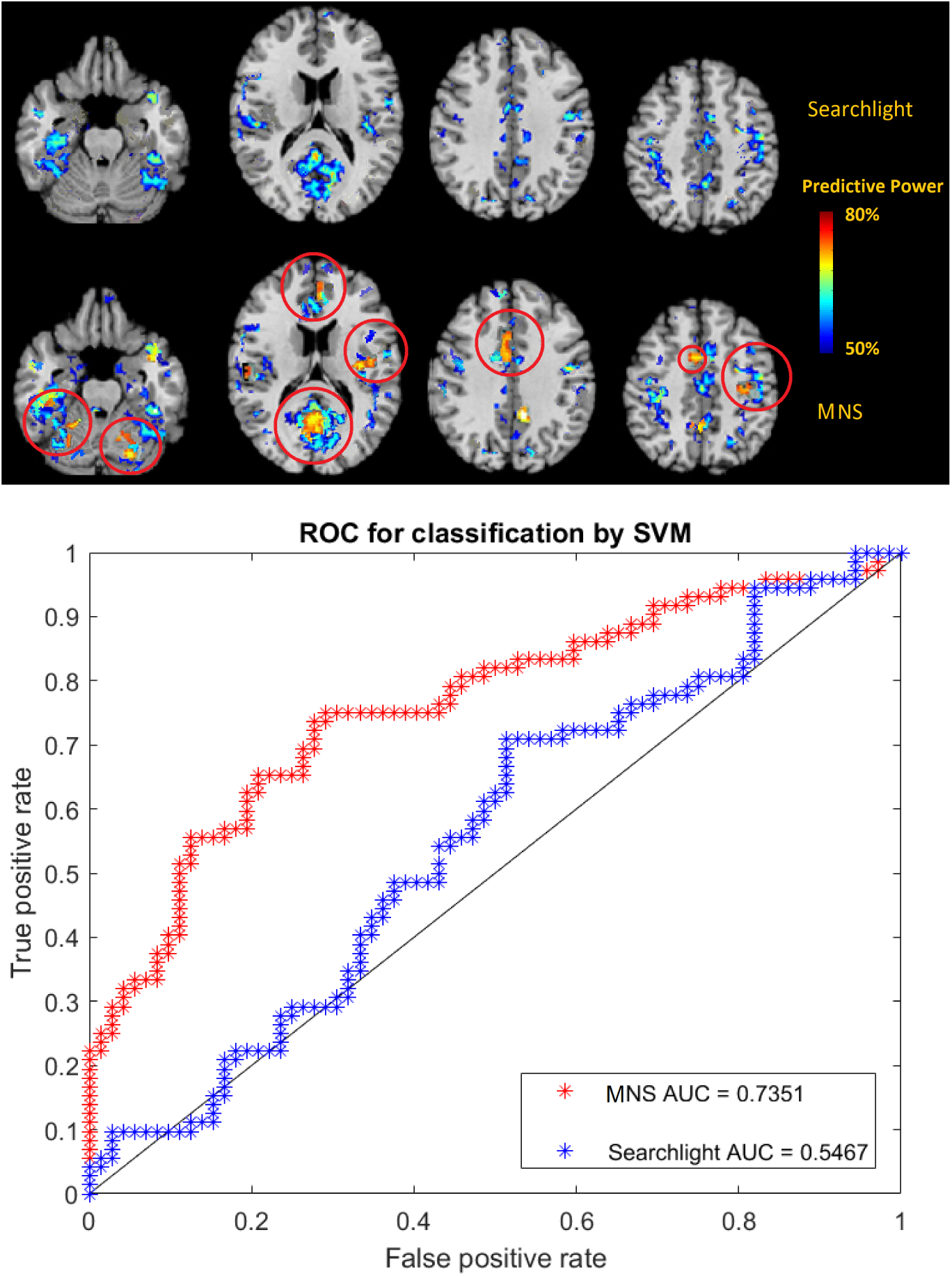
Top: A comparison of the above chance accuracy clusters derived using the searchlight criteria (top row) and the MNS algorithm (bottom row) on the ABIDE data set. Major differences between the two maps are indicated by the red circles. In case of overlapping clusters generated by MNS, the clusters with the highest predictability were selected for this visualization. Bottom: Classification performance on full-brain search space for ABIDE dataset based on the above chance clusters as the features where the train-test population was 546-137 respectively.

**Figure 5.**
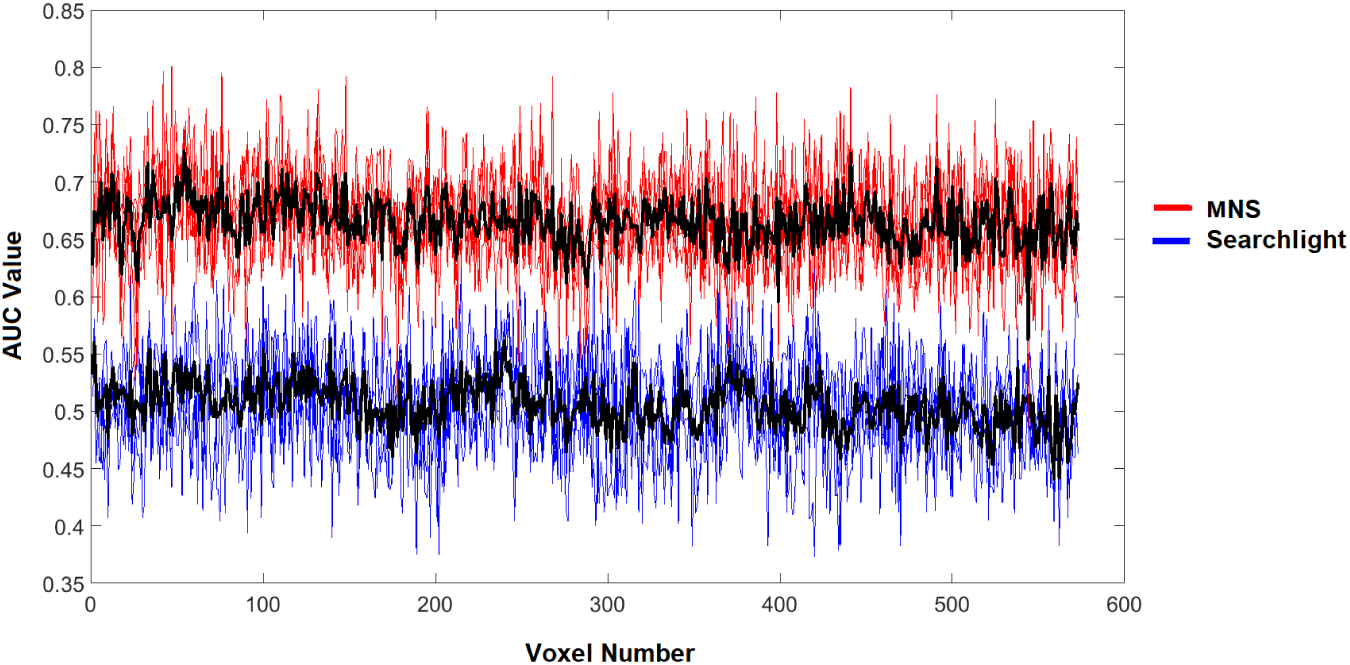
Comparison of classification AUC with svm between MNS and Searchlight with 3 voxel radius for left crus II of the cerebellum (region 93 per AAL) with 573 voxels.

**Figure 6.**
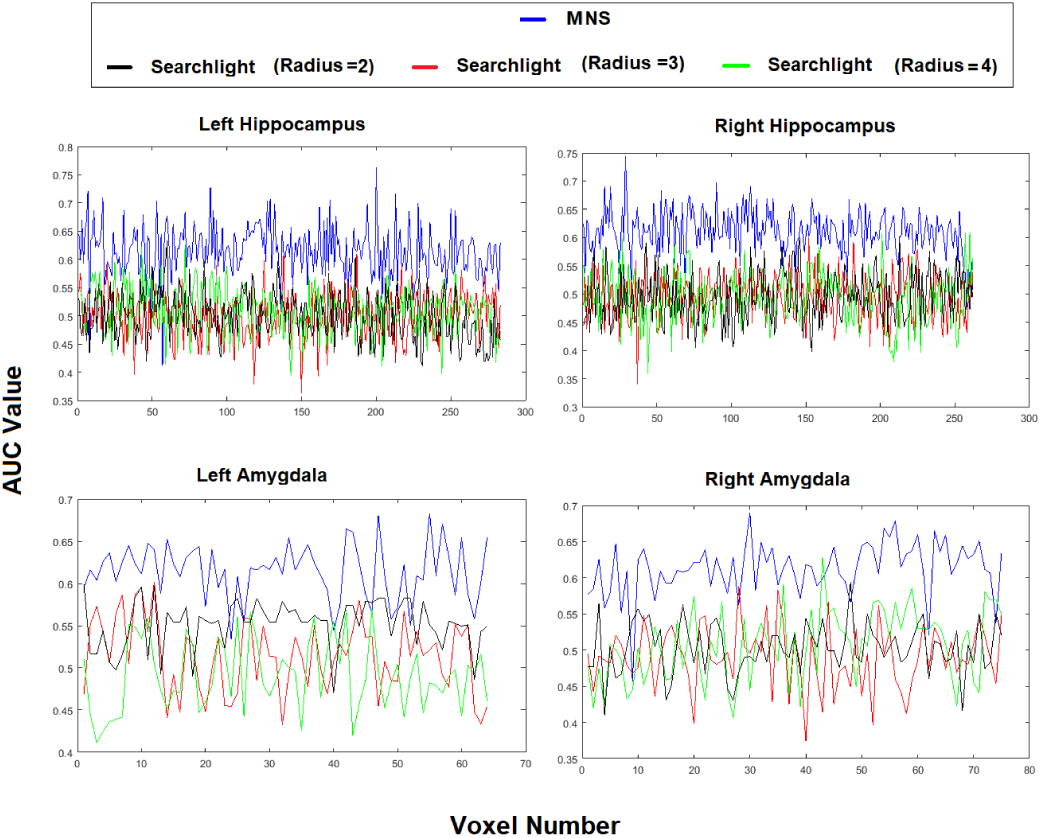
Test AUC for classification with svm based on the MNS algorithm and the searchlight method with different search radii for right and left hippocampus and amygdala from the ABIDE dataset.

**Figure 7.**
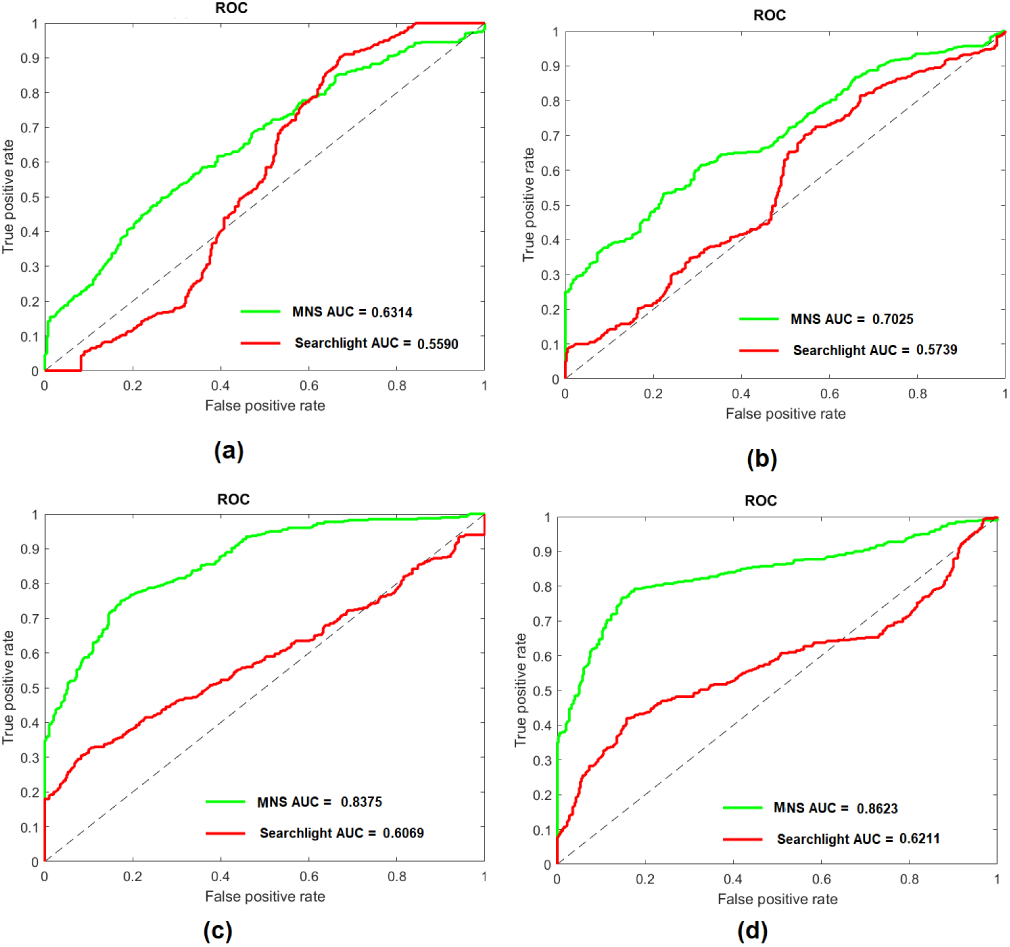
Area under the ROC curve for synthetic datasets of size 100 (a), 500 (b), 10,000 (c), and 30,000 (c) voxels.

## Discussion

We suggested a new method for MVPA analysis with the objective of increasing the precision of the voxel-level information map while eliminating several constraints including the requirement for parameter tuning. We proposed a data-driven approach which performs a search based on a data-driven heuristic function, and starting from each voxel, dynamically detects the appropriate formation of its combination with other voxels in its vicinity. This approach presented several valuable advantages over the other decoding methods, which we discuss in this section.

### Information of voxel clusters are discovered precisely

Due to data-driven expansion of clusters around each starting voxel, their calculated information is automatically assigned to the individual clusters without redundant voxels. Therefore, precise knowledge about the discriminant power of clusters is provided by avoiding exaggeration of the spatial boundaries of informative areas. This property removes the possibility of discontinuous detection, i.e. existence of one or few highly informative voxels dominating the information of a region and increasing the possibility of overfitting.

### Parameter tuning is not required

As described in the methodology section, the only input to the MNS algorithm are the sets of time series of two or more populations of subjects averaged over time, i.e. the BOLD activation distribution of two or more populations. The elimination of tuning parameters such as searchlight shape (spherical or cubical) and radius, or machine learning hyper parameters (example: deep learning based methods) or regularization parameters (the *λ* value in LASSO or elastic net) not only increases the generalizability of the results, but also increases the efficiency of analysis. The latter point is due to the fact that the requirement for multiple runs of the analysis with different searchlight radii and then selecting the highest performing parameters is removed. Moreover, this property of the MNS algorithm resolves the issue of heterogeneous accuracy on various regions which normally occurs due to assignment of a single searchlight radius for larger brain regions with various topological characteristics. Plus, parameter tuning requires researchers to have advanced expertise of the methods they intend to use.

### The shape of the clusters are not bound by any constraints

The shape of the searchlight sphere can affect the information detection precision. For example, in the presence of an elliptical cluster, a spherical searchlight could fail to detect its complete boundaries, thus creating an imprecise information map. However, the shape of the clusters created by MNS merely depends on the information of the voxels and their combination with the voxels in their vicinity. This can specifically be a useful property for clusters located at the edges of the search space that are more prone to irregular shapes.

### Prediction accuracy is enhanced simultaneous to cluster detection

The search space traversal of MNS is navigated toward voxels that increase the information of the clusters as its objective is to solve the optimization function that finds the combination of neighboring voxels that maximize the information. Therefore, starting from each voxel, the search continues as long as possibility exists for enhancing the discrimination power of the cluster by adding useful voxels and removing redundant ones. As discussed in the results section, to validate the generalization of the selected feature subsets, the classification accuracy can be calculated through ground truth cross-validation.

### Additional analysis is not required for interpretation of the information map

Due to data-driven assignment of discriminant scores on the voxel level, clusters with clear boundaries are generated by the MNS method. Therefore, unlike some of the other MVPA methods, complementary tests are not required to detect informative voxels within the clusters. Moreover, the proposed approach provides higher intuitiveness compared to methodologies often considered as “black box” such as deep learning based approaches or regularization-based methods.

### Global optimality

The conventional step-wise greedy search method for feature selection yields suboptimal feature subsets due to falling in local minima ^50^. This is due to the fact that the choice of features depends on the order of their selection. Therefore, the final feature set is not necessarily the globally optimal set. However, in the approach proposed in this paper, for every starting voxel, multiple useful neighbors are admitted at each neighborhood layer (step 2 of the algorithm), therefore reducing the chance of falling in local optima. Moreover, the output of the algorithm includes the discovered information clusters starting from every voxel in the search space. In other words, steps 1 to 4 are performed for every voxel in the search space, resulting in a more thorough search over possible voxel combinations. While this algorithm does not perform an exhaustive search over every possible combination of voxels, these two properties significantly decrease its chance of falling into local optima, therefore providing a set of solutions that makes it near optimal. Moreover, global information within clusters are considered through both spectral relevance analysis and the redundancy analysis as mentioned before during each step of the search and reevaluation of the information cluster. Note that the problem of finding the best set of features is a significantly more complex problem than distance-based graph search approaches. This is due to non-linear relations between the features compared with the notion of physical distance which can be measured by accumulating the subdistances in the search space.

### Time Complexity Analysis

A brute force implementation of the proposed search algorithm would require multiple calculations for every attribute to be performed repeatedly due to the overlap among information clusters. However, by exploiting these overlaps, we proposed a dynamic programming implementation of the calculations where each of theses calculations only takes place once for each voxel in the search space. In case of revisiting voxels, instead of repeating the calculations, previous calculations stored in hash tables are looked up, thus increasing the efficiency of the algorithm significantly. The majority of calculation of the spectral relevance analysis is also calculated prior to search by avoiding repetition of the matrix multiplications in equation 9. Time complexity of spectral intra-group selection with *m* dimensions (in our case, voxels) is *O*(*m*). Another point worth mentioning is that the redundancy analysis is only performed after the set of high quality neighbors at each proximity layer are admitted to the feature vector, which facilitates a more rapid traversal in the search space. The worst case scenario happens when the algorithm visits every voxel for discovering each information cluster, meaning that the entire search space increases the information of the initial cluster. However, in practice this is a rare case. In fact, our empirical results showed that this algorithm traverses a much smaller subspace of the search space, which reduces its time complexity. Nevertheless, since the algorithm is completely data-driven, the time that the proposed procedure takes to run on a given dataset highly depends on the data itself. However, the algorithms run time in the heaviest experimental setup for our experiment, which was the whole-brain analysis for the entire dataset, was conducted within few hours, which is within the norm of feasible analysis.

## Materials and Methods

The pseudo code for the MNS algorithm is provided in Algorithm 1. Here we explain the steps of the proposed methodology in more detail. As can be seen in that pseudo code, the input to the proposed approach includes an *M* × *P* matrix *X* where *M* is the number of subjects and *P* is the number of search space voxels, and a vector *Y* containing the labels. Therefore, each element *X_ij_* contains the averaged time course of voxel *j* for subject *i* over time. Note that the input to this algorithm can include more than two groups, and similar setup can be designed for multi-class scenario. As the search starts, the spectral discriminant score of Cluster *C_temp_*, which initially only includes the voxel *V_start_* is measured by exploiting the pre-calculated local and global affinity matrix multiplications based on the right side component of equation 9(line 10). Then, for each neighboring voxel of *C_temp_*, the spectral discriminant score of the addition of the neighbors *V_neighbor_* and *C_temp_* are calculate and compared with the score of *C_temp_*, and useful voxels are added to *C_temp_* to expand it (lines 11 to 22). After one step of neighborhood search and expansion of *C_temp_*, redundancy criteria is performed by first looking up the matrices that will contain the calculations for three components of equation 7 for each voxel. If the corresponding values for the voxels inside *C_temp_* do exist in the vector *Feature_inf_* and matrices *Interact_u_* and *Interact_c_*, (meaning that the voxel has been previously visited), the values are directly used to avoid recalculations. Otherwise, the mutual information elements are calculated and placed in its corresponding location in the three variables for future use (lines 23 to 30).

The redundancy scores of each voxel inside *C_temp_* is then compared with a pre-specified threshold *T* to remove redundant features (lines 31 to 33). Note that *T* can be assigned in various fashions, depending on how strict the redundancy analysis is preferred to be. Note that in this algorithm we avoid removing the recently added voxels during the redundancy step to avoid the possibility of falling into infinite loops by repeatedly adding and removing the same voxels (note the loop condition in line 23 which excludes the newly added voxels). For the same reason, we avoid adding newly removed voxels (note the condition in line 12). If the algorithm does not find any more useful neighbors during the search, after saving the results in the output it moves to the voxel next to *V_start_*, and pursues the same steps until it covers the entire search space, providing a complete list of information clusters and their information value (lines 36 to 43).

In the next sections we discuss the spectral discriminant analysis and interaction information based redundancy analysis in more detail.

### Data Preparation

Since the MNS performs a search among neighbors of clusters, it is necessary to create the neighborhood matrix of the voxels before starting the search procedure, and just point the algorithm to the neighbors of the voxel during the search process. The neighborhood matrix can be created by simply measuring the euclidean distance of the voxels based on their three dimensional coordinates. In this matrix, called matrix *M_neighbors_*, each row belongs to a voxel, and the columns include the indices of their immediate neighboring voxels. Note that topological properties of specific regions can be considered during pre-processing using FMRIB Software Library (FSL), and then the neighborhood matrix can be created. This would not affect the performance of the search, as it only requires the indices of the neighboring voxels.

#### Spectral Discriminant Analysis Via Graph Laplacian

In order to admit a newly arrived voxel *v_i_* to the information cluster *C_v_* in our online feature analysis scheme, the relevance of its addition to the cluster is compared to the relevance of the cluster before its arrival. Most feature evaluation criteria measure the quality of the attributes individually. Also, many feature set evaluation methods perform a linear evaluation of them. However, linear methods fail if the data lies on a low-dimensional manifold because the data structure becomes highly nonlinear. Therefore, we use non-linear approaches for both relevance and redundancy analysis. As the relevance measurement, we employ spectral graph clustering approach which takes the geometric and topological properties of a given manifold into account.

The spectral graph theory for feature selection attempts to find a smooth feature selector matrix based on the notions of class affiliation which measures the ratio between local and global affinity. In other word, in higher the following relation, the higher quality is the feature.

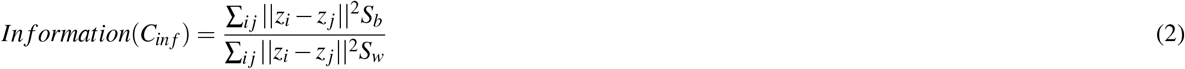

Where *Z* is a transformation of *X* by the feature space projection *Z* = *W^T^X*, and *z_i_* and *z_j_* are the corresponding values of *z* for data points *i* and *j*. Also, *S_b_* and *S_w_* denote the Fisher score between and within class adjacency matrices ^39^ which are calculated as below:

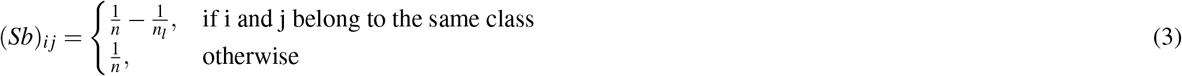

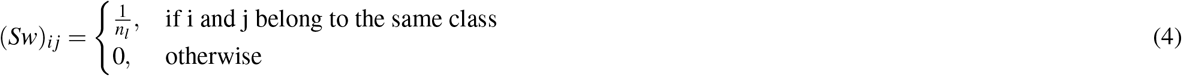

Where *n_l_* denotes the number of data points from class *l*. Given the adjacency matrix *S_w_* and the degree matrix *D_w_* of the graph *G_w_* being defined as *D_w_* = *diag*(*S*_*w*1_) if *i* = *j*, and 0 otherwise, the Laplacian matrix of graph *G_w_* (within distance) is defined as *L*_*w*=_*D_w_* – *S_w_* ^51^. Similarly, the Laplacian matrix of graph *G_b_* (between distance) is defined as *L_b_* = *D_b_* – *S_b_*.

##### Algorithm 1 MRMR Neighborhood search with Dynamic Programming

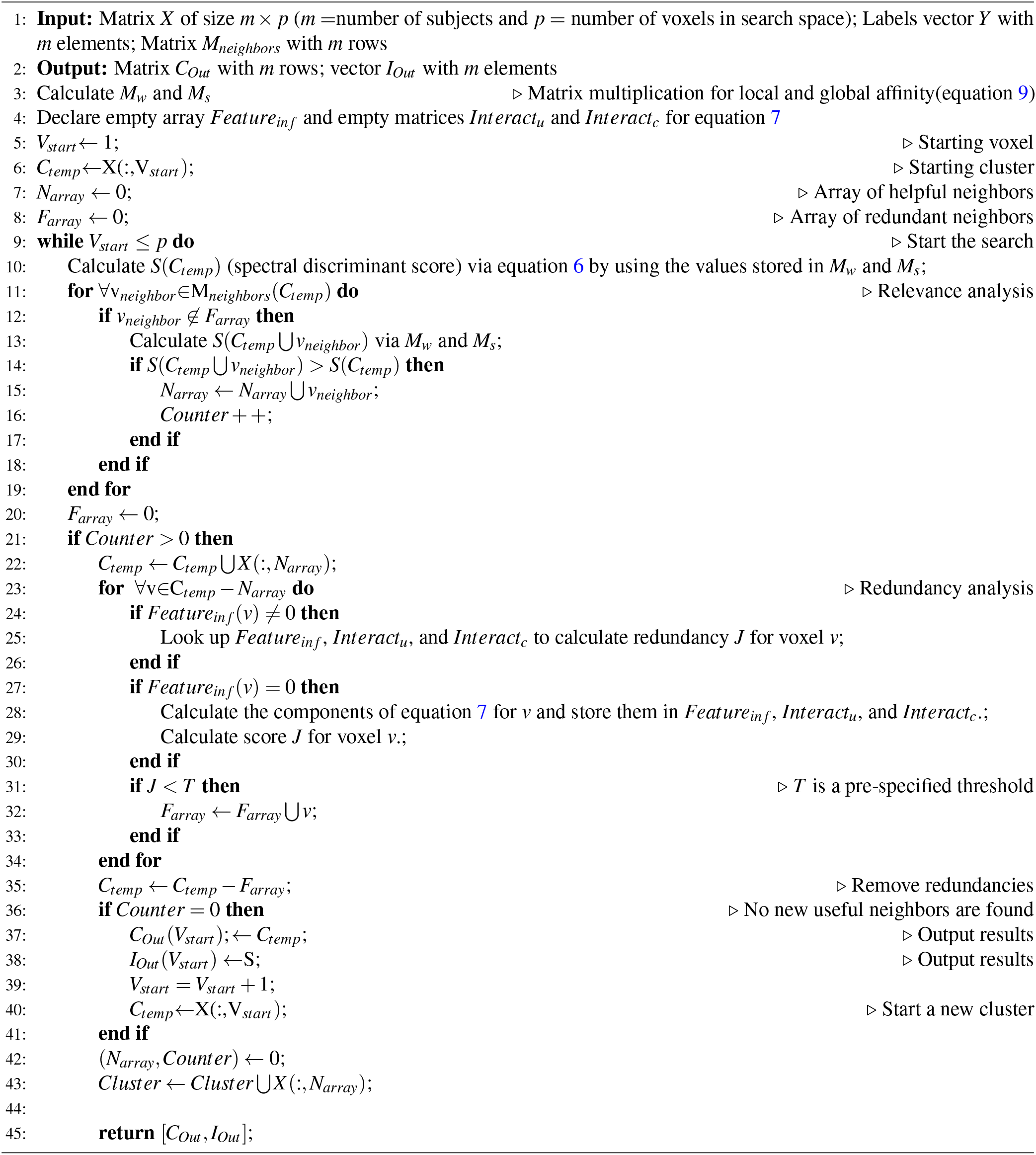

Moreover, with the procedure of feature selection, data matrix *X* is transformed to *Z* ∈ *R*^*m*×*n*^ by the feature space projection *Z* = *W^T^* × *X*. This criteria can be converted into the trace ratio of the form below based on the Laplacian matrix of the graph *G*(*C_inf_*)^38, 39^:

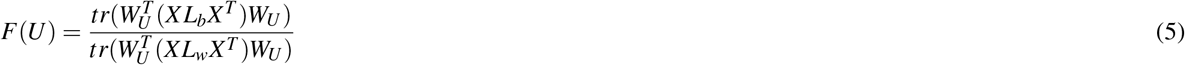

And the spectral quality for the arriving feature *f_i_* can be measured by a score defined below:

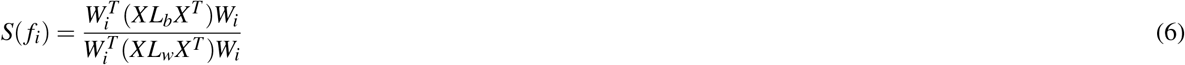

### Inter-group feature redundancy analysis with spatial bias

Here we introduce the inter-group feature set analysis which aims to obtain the optimal subset. For this purpose, three mutual information elements are calculated to consider both the global and local interactions within the set of voxels which is formulated below ^42^.

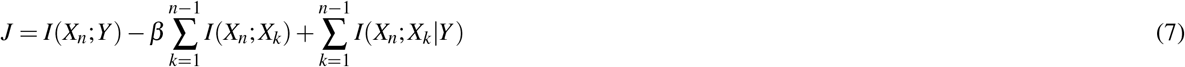

Where *I* denotes the mutual information between two vectors, *X* is the set of features, and *Y* denotes the labels, and *n* is the number of features. The first term is the mutual information between the individual feature and the class label, the second term calculates the sum of mutual information between each feature and every other feature in the set, and the third term (rightmost) calculates the interaction between the features at the conditional presence of the target feature. The variables *β* penalizes high correlations between the features to further signify the redundant features. As mentioned previously, we introduce a bias based on the spatial location of the voxels with regards to the clusters in the redundancy analysis. Therefore, the term *β* is assigned as the inverse degree centrality of the voxel wih regards to the information cluster. This constraint also removes the necessity of tuning the parameter *β* as it is automatically calculated by the algorithm. As mentioned previously, by punishing redundancies at the fringes of the clusters, this spatial bias alleviates the issue of noise in voxel level resolution by considering the smooth proximal neural activation patterns. Also, the mutual information *I* is calculated by the formula below:

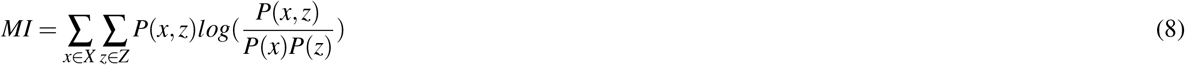

Where *x* denotes the feature values and *z* shows the class labels.

### Dynamic Programming-based Implementation

We suggest a top-down dynamic programming approach for the proposed method to increase its run time efficiency by exploiting the existing overlapping subproblems in the search space. This approach is performed for both relevance and redundancy analysis criteria. For relevance criteria, based on equation 6, the majority of computational burden is on the matrix multiplication *XL_b_XT* which is derived from the equation below based on the property of Laplacian matrix (Note that similar property exists for *S_b_*):

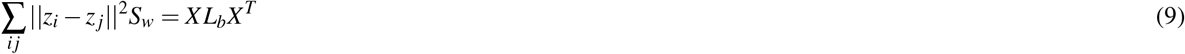

However, we can make the above matrix multiplication only once, before starting the search, and preserve the values inside a hash table instead of recalculating it during search. Due to the property of trace of matrices, the value for each voxel can be easily obtained by looking up the pre-calculated hash table.

Moreover, for redundancy analysis, we break down the calculations of each voxel for each of the three elements in equation 7. This is due to the fact that both the unconditional and conditional mutual interaction informations between each voxel and every other voxel is a sum calculation, and the individual value of each feature(the leftmost element) can be looked up after being calculated and stored once. Therefore, each time redundancy analysis is being performed on a feature set *C_v_*, for each voxel, the algorithm first checks if the calculation for it exists in the hash matrices, and only performs the calculations and saves them if they have not been performed previously. This criteria is displayed in Algorithm 1. Note that the calculations for 7 based on this approach are only performed when needed, which is more efficient than pre-calculating the interactions between each voxel and every other voxel in the search space. The value of the global interaction elements in *J* score according to equation 7 are calculated recursively using previous calculations. This is due to the fact that based on the proposed search method, we are interested in detecting redundancies within an analytically-formed cluster of voxels which only requires the interaction of voxel being visited with the members of the cluster rather than its interaction with every other voxel in the entire search space. As mentioned previosuly, this dynamic programming based implementation removes the requirement of repeating calculations by exploiting overlapping subproblems in the search.

## Supporting information

Supplemental Results

## References

1. Huettel, S. A., Song, A. W., McCarthy, G. et al. Functional magnetic resonance imaging, vol. 1 (Sinauer Associates Sunderland, MA, 2004).

2. Wong, A. C.-N., Palmeri, T. J., Rogers, B. P., Gore, J. C. & Gauthier, I. Beyond shape: how you learn about objects affects how they are represented in visual cortex. PloS one 4, e8405 (2009).

3. Norman, K. A., Polyn, S. M., Detre, G. J. & Haxby, J. V. Beyond mind-reading: multi-voxel pattern analysis of fmri data. Trends cognitive sciences 10, 424–430 (2006).

4. Davis, T. et al. What do differences between multi-voxel and univariate analysis mean? how subject-, voxel-, and trial-level variance impact fmri analysis. Neuroimage 97, 271–283 (2014).

5. Swearingen, J. Characterizing the temporal dynamics in functional connectivity measured with fMRI (Medical University of South Carolina, 2015).

6. Jimura, K. & Poldrack, R. A. Analyses of regional-average activation and multivoxel pattern information tell complementary stories. Neuropsychologia 50, 544–552 (2012).

7. Gardumi, A. et al. The effect of spatial resolution on decoding accuracy in fmri multivariate pattern analysis. Neuroimage 132, 32–42 (2016).

8. Stelzer, J., Chen, Y. & Turner, R. Statistical inference and multiple testing correction in classification-based multi-voxel pattern analysis (mvpa): random permutations and cluster size control. Neuroimage 65, 69–82 (2013).

9. Etzel, J. A., Zacks, J. M. & Braver, T. S. Searchlight analysis: promise, pitfalls, and potential. Neuroimage 78, 261–269 (2013).

10. Kriegeskorte, N., Goebel, R. & Bandettini, P. Information-based functional brain mapping. Proc. Natl. Acad. Sci. 103, 3863–3868 (2006).

11. Kriegeskorte, N. & Bandettini, P. Analyzing for information, not activation, to exploit high-resolution fmri. Neuroimage 38, 649–662 (2007).

12. Uddin, L. Q. et al. Multivariate searchlight classification of structural magnetic resonance imaging in children and adolescents with autism. Biol. psychiatry 70, 833–841 (2011).

13. Chen, Y. et al. Cortical surface-based searchlight decoding. Neuroimage 56, 582–592 (2011).

14. Shimizu, Y. et al. Toward probabilistic diagnosis and understanding of depression based on functional mri data analysis with logistic group lasso. PloS one 10, e0123524 (2015).

15. Toiviainen, P., Alluri, V., Brattico, E., Wallentin, M. & Vuust, P. Capturing the musical brain with lasso: Dynamic decoding of musical features from fmri data. Neuroimage 88, 170–180 (2014).

16. Ng, B. & Abugharbieh, R. Generalized sparse regularization with application to fmri brain decoding. In Biennial International Conference on Information Processing in Medical Imaging, 612–623 (Springer, 2011).

17. Gramfort, A., Thirion, B. & Varoquaux, G. Identifying predictive regions from fmri with tv-l1 prior. In Pattern Recognition in Neuroimaging (PRNI), 2013 International Workshop on, 17–20 (IEEE, 2013).

18. Tibshirani, R., Saunders, M., Rosset, S., Zhu, J. & Knight, K. Sparsity and smoothness via the fused lasso. J. Royal Stat. Soc. Ser. B (Statistical Methodol. 67, 91–108 (2005).

19. Wolz, R., Aljabar, P., Hajnal, J. V. & Rueckert, D. Manifold learning for biomarker discovery in mr imaging. In International Workshop on Machine Learning in Medical Imaging, 116–123 (Springer, 2010).

20. Belkin, M. & Niyogi, P. Laplacian eigenmaps for dimensionality reduction and data representation. Neural computation 15, 1373–1396 (2003).

21. Dechter, R. & Pearl, J. Generalized best-first search strategies and the optimality of a. J. ACM (JACM) 32, 505–536 (1985).

22. DeVore, R. A. & Temlyakov, V. N. Some remarks on greedy algorithms. Adv. computational Math. 5, 173–187 (1996).

23. Dijkstra, E. W. A note on two problems in connexion with graphs. Numer. mathematik 1, 269–271 (1959).

24. Prim, R. C. Shortest connection networks and some generalizations. Bell system technical journal 36, 1389–1401 (1957).

25. Hart, P. E., Nilsson, N. J. & Raphael, B. A formal basis for the heuristic determination of minimum cost paths. IEEE transactions on Syst. Sci. Cybern. 4, 100–107 (1968).

26. Burke, E. K. et al. Hyper-heuristics: A survey of the state of the art. J. Oper. Res. Soc. 64, 1695–1724 (2013).

27. Mladenovic, N. & Hansen, P. Variable neighborhood search. Comput. & operations research 24, 1097–1100 (1997).

28. Tan, G.-z., He, H. & Aaron, S. Global optimal path planning for mobile robot based on improved dijkstra algorithm and ant system algorithm. J. Cent. South Univ. Technol. 13, 80–86 (2006).

29. Preparata, F. P. & Shamos, M. I. Computational geometry: an introduction (Springer Science & Business Media, 2012).

30. Balakrishnama, S. & Ganapathiraju, A. Linear discriminant analysis-a brief tutorial. Inst. for Signal information Process. 18, 1–8 (1998).

31. Welling, M. Fisher linear discriminant analysis. Dep. Comput. Sci. Univ. Tor. 3 (2005).

32. Wilks, D. S. Cluster analysis. In International geophysics, vol. 100, 603–616 (Elsevier, 2011).

33. Rousseeuw, P. J. Silhouettes: a graphical aid to the interpretation and validation of cluster analysis. J. computational applied mathematics 20, 53–65 (1987).

34. Zhao, Z. & Liu, H. Spectral feature selection for supervised and unsupervised learning. In Proceedings of the 24th international conference on Machine learning, 1151–1157 (ACM, 2007).

35. Perkins, S. & Theiler, J. Online feature selection using grafting. In Proceedings of the 20th International Conference on Machine Learning (ICML-03), 592–599 (2003).

36. Zhou, J., Foster, D., Stine, R. & Ungar, L. Streaming feature selection using alpha-investing. In Proceedings of the eleventh ACM SIGKDD international conference on Knowledge discovery in data mining, 384–393 (ACM, 2005).

37. Wu, X., Yu, K., Ding, W., Wang, H. & Zhu, X. Online feature selection with streaming features. IEEE transactions on pattern analysis machine intelligence 35, 1178–1192 (2013).

38. Wang, J. et al. Online feature selection with group structure analysis. IEEE Transactions on Knowl. Data Eng. 27, 3029–3041 (2015).

39. Nie, F., Xiang, S., Jia, Y., Zhang, C. & Yan, S. Trace ratio criterion for feature selection. In AAAI, vol. 2, 671–676 (2008).

40. Koller, D. & Sahami, M. Toward optimal feature selection. Tech. Rep., Stanford InfoLab (1996).

41. Aliferis, C. F., Statnikov, A., Tsamardinos, I., Mani, S. & Koutsoukos, X. D. Local causal and markov blanket induction for causal discovery and feature selection for classification part i: Algorithms and empirical evaluation. J. Mach. Learn. Res. 11, 171–234 (2010).

42. Brown, G. A new perspective for information theoretic feature selection. In Artificial intelligence and statistics, 49–56 (2009).

43. Bellman, R. Dynamic programming (Courier Corporation, 2013).

44. Bellman, R. E. & Dreyfus, S. E. Applied dynamic programming, vol. 2050 (Princeton university press, 2015).

45. Di Martino, A. et al. The autism brain imaging data exchange: towards a large-scale evaluation of the intrinsic brain architecture in autism. Mol. psychiatry 19, 659 (2014).

46. Baron-Cohen, S. et al. The amygdala theory of autism. Neurosci. & Biobehav. Rev. 24, 355–364 (2000).

47. Schumann, C. M. et al. The amygdala is enlarged in children but not adolescents with autism; the hippocampus is enlarged at all ages. J. neuroscience 24, 6392–6401 (2004).

48. Bauman, M. & Kemper, T. L. Histoanatomic observations of the brain in early infantile autism. Neurology 35, 866–866 (1985).

49. Fatemi, S. H. et al. Consensus paper: pathological role of the cerebellum in autism. The Cerebellum 11, 777–807 (2012).

50. Vafaie, H. & Imam, I. F. Feature selection methods: genetic algorithms vs. greedy-like search. In Proceedings of the International Conference on Fuzzy and Intelligent Control Systems, vol. 51, 28 (1994).

51. Mohar, B., Alavi, Y., Chartrand, G. & Oellermann, O. The laplacian spectrum of graphs. Graph theory, combinatorics, applications 2, 12 (1991).

