## Supplemental Results for "A heuristic feature cluster search algorithm for precise functional brain mapping"

### Supporting information

January 12, 2019

#### Comparison of Prediction Results

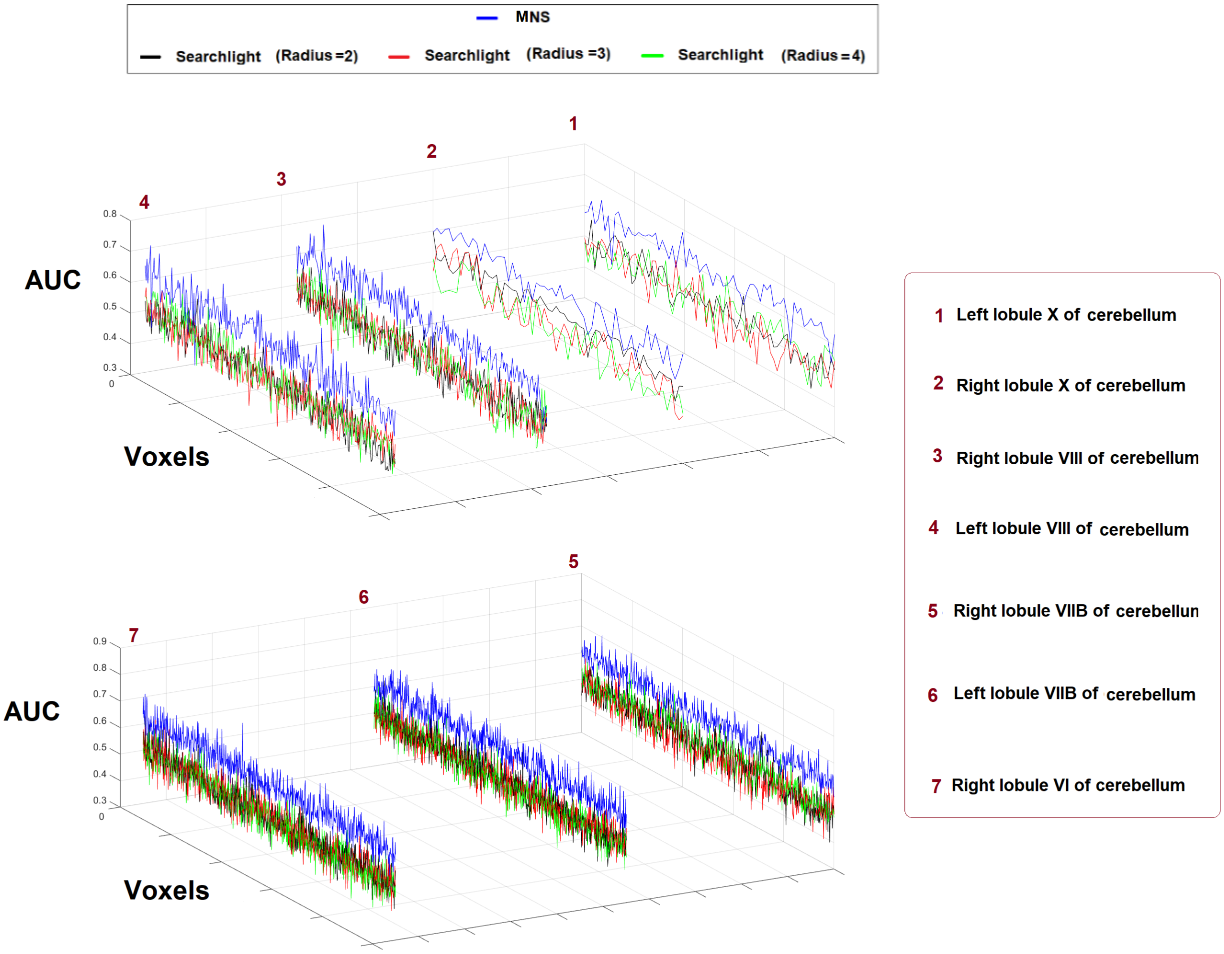


Figure 1: Test AUC for classification with decision tree of the MNS algorithm and the searchlight method with different search radii for different cerebellar regions per AAL using the ABIDE dataset.
